# Forebrain-specific loss of erythropoietin provokes compensatory upregulation of different EPO receptors

**DOI:** 10.1101/2025.09.11.675614

**Authors:** Umer Javed Butt, Umut Çakır, Anne-Fleur Wildenburg, Yasmina Curto, Liu Ye, Vikas Bansal, Susann Boretius, Klaus-Armin Nave, Manvendra Singh, Hannelore Ehrenreich

## Abstract

The procognitive growth factor erythropoietin (EPO) and its canonical receptor, EPOR, have long been recognized to be expressed by most cell types in the brain. Cognitive domains, improved by injections of exogenous EPO or by endogenous, hypoxia-stimulated EPO, include important forebrain functions, namely attention, working memory, drive, and executive performance. To gain mechanistic insight into the involvement of forebrain-expressed EPO, we deleted EPO in mice using as specific cre-driver *Emx1*. Here, we report that these mutant mice act comparably to their wildtype littermates in a comprehensive behavioral test battery. Importantly, we find that the transcripts of both EPOR and a novel, brain-expressed EPO receptor, EphB4, respond to EPO deletion with compensatory upregulation. EphB4 expression in brain and its increase upon forebrain erasure of EPOR are confirmed by *in situ* hybridization and immunohistochemistry. The augmented expression of both EPOR and EphB4 and their regulatory intercorrelation may explain why *EmxEPO* mutants show an even superior performance in the most challenging working memory task. Using the previously published single-nuclei-RNA-seq dataset, we further confirm the suggested compensatory mechanism, wherein EPO loss or reduction drives elevated EPOR expression, adding another layer to the intricate regulation of EPO signaling in hippocampal pyramidal neurons. Collectively, these data may explain the lack of behavioral and negative cognitive consequences upon forebrain-wide EPO elimination.

## INTRODUCTION

Mammalian forebrain tasks comprise all facets of higher cognition, including executive functions, processing speed, attention, learning and memory, complex multisensory networking as well as drive, motivation and emotions [1–3]. Recombinant human (rh) erythropoietin (EPO), applied intravenously, as well as endogenous, brain-expressed EPO have been recognized for decades to exert a substantial modulating influence on these core competencies of the central nervous system [4]. We identified pivotal procognitive roles of hypoxia-induced EPO in brain, which are imitated by rhEPO treatment. These roles are part of a fundamental regulatory circle, in which neuronal networks - challenged by motor-cognitive tasks - drift into transient hypoxia, thereby triggering neuronal EPO and EPO receptor (EPOR) expression [4, 5]. In fact, complex motor-cognitive exercise causes hypoxia across essentially all brain areas, with hypoxic neurons particularly abundant in the hippocampus [6, 7]. Conducting transcriptional hippocampal profiling of rhEPO-treated mice, we discovered populations of newly differentiating pyramidal neurons, overpopulating to ∼200% upon rhEPO with upregulation of genes crucial for neurodifferentiation, dendrite growth, synaptogenesis, memory formation, and cognition [8].

An alternative EPO binding site, EphB4, had originally been reported in tumor cells as a member of the family of Ephs, i.e. EPO-producing human hepatocellular receptors [9–11]. Modulation of NMDA receptor-dependent calcium influx and gene expression, as shown for other EphB receptors [12], could be among the physiological functions also of EphB4 receptors in brain, but remains to be demonstrated. EphB4, on the other hand, seems to be indirectly involved in adult hippocampal neurogenesis [13] and to play an essential part in regulating small artery contractility and blood pressure [14].

In order to get deeper insight into the physiological significance of EPO and its receptors, we generated and studied mice lacking EPO expression in the forebrain. Here, we report that this lack is apparently compensated for by upregulation of EPOR and the novel EPO receptor in brain, EphB4. Moreover, we provide first evidence that regulation of EPOR and EphB4 are interrelated.

## METHODS

### Mice

All experiments were conducted by investigators unaware of genotypes and group assignment (′fully blinded′) and in accordance with the local authorities (Animal Care and Use Committee: Niedersächsisches Landesamt für Verbraucherschutz und Lebensmittelsicherheit, LAVES) following the German Animal Protection Law (AZ 33.19-42502-04-18/2803 & AZ 33.19-42502-04-17/2393). Mice were segregated based on sex and genotype and kept in group-housing environment in type IV cages (Techniplast Hohenpeiβenberg, Germany) inside ventilated cabinets (Scantainers, Scanbur Karlsunde, Denmark) unless stated otherwise. The cages were furnished with wood-chip bedding and nesting material (Sizzle Nest, Datesand, Bredbury, United Kingdom). All mice were housed in the animal facility of the Max Planck Institute for Multidisciplinary Sciences in a temperature-controlled environment (21±2°C, humidity ∼50%) on a 12h light/dark cycle. They had access to food (Sniff Spezialdiäten, Bad Soderberg, Germany) and water *ad libitum,* except for the female *EmxCre::EPO* mice during the IntelliCage behavioral testing paradigm (see below).

### Mouse model and genotyping

To delete EPO expression in forebrain, we crossed mice in which *EPO* exons 2-3 are flanked by loxP-sites with a Emx1IREScre driver line under control of *Emx1* promoter [15] (**Figure 1A**). Emx1IREScre refers to the knock-in allele in which a bicistronic IRES-Cre recombinase cassette has been inserted downstream of the endogenous Emx1 coding sequence. EmxIREScre^+/+^ and EmxIREScre^+/-^ designates mice, homozygous (WT) and heterozygous (KO) for Emx1IREScre and therefore controlling activity of cre recombinase. EmxEPO mice were maintained on C57BL/6N background. Experimental groups for basic behavior battery included female EPO*^flox/flox*^*Emx*^IREScre+/+^* (EPO WT) and female EPO*^flox/flox*^*Emx*^IREScre+/-^* (EPO KO).

**Figure 1:**
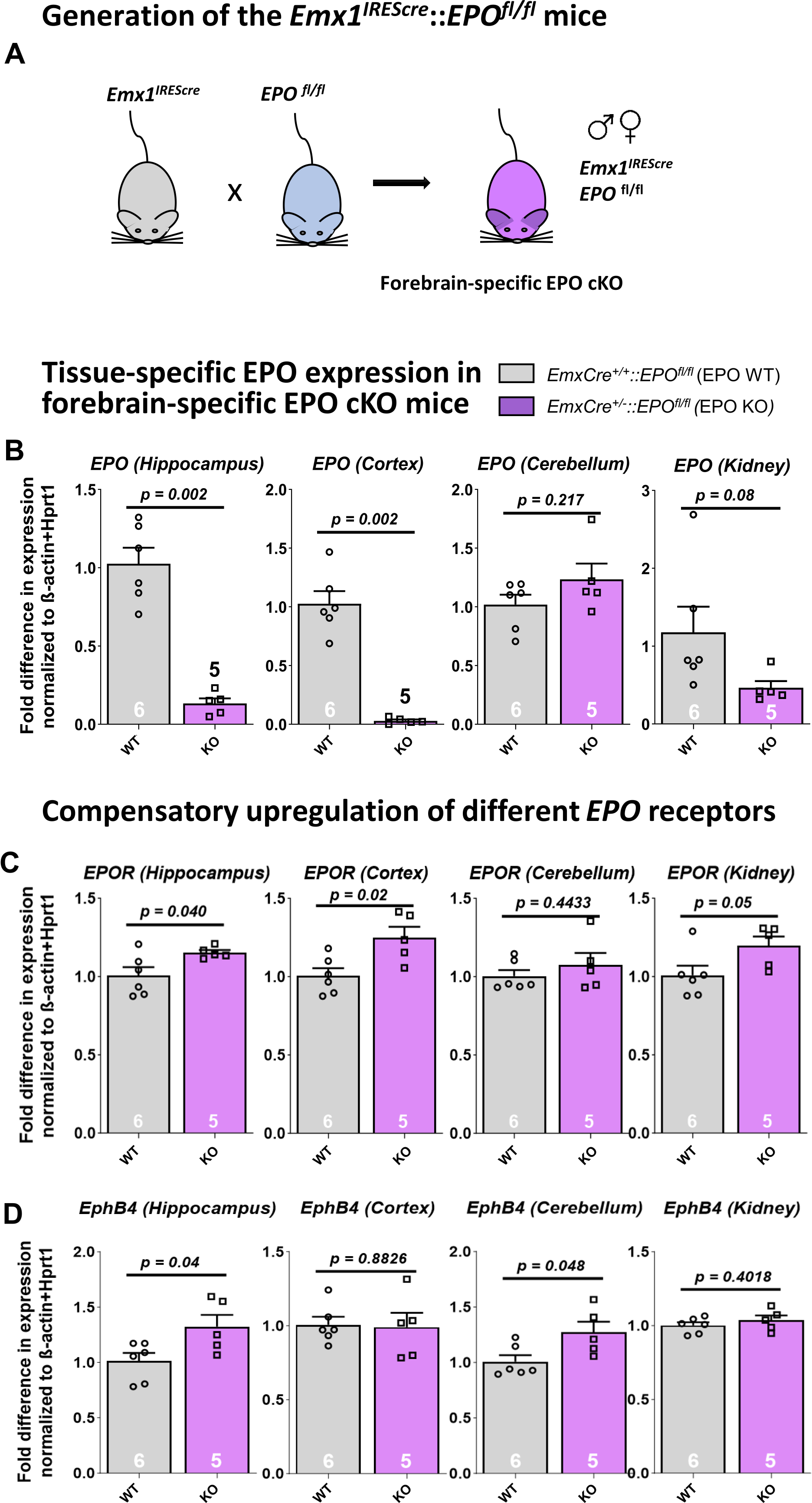
Generation of forebrain-specific *EPO* KO mice and tissue-specific expression of EPO and EPO receptors. **(A)** Schematic representation of the generation of the *EmxEPO* KO mouse line. Forebrain-specific loss of *EPO* was achieved by crossing the *Emx1^IREScre^* driver line with mice carrying a floxed *EPO* gene. **(B)** *EPO* mRNA expression (normalized to *β-actin* and *Hprt1*) in hippocampus, cortex, cerebellum and kidney of *EPO* KO and WT control mice. **(C-D)** *EPOR and EphB4* mRNA expression (normalized to *β-actin* and *Hprt1*) in hippocampus, cortex, cerebellum and kidney of *EPO* KO and WT control mice. All data are shown as mean±SEM with p<0.05 considered significant. Statistical analyses were conducted via unpaired t-tests with Welch’s correction.

Forebrain-specific loss of EPO did not result in any overt effect on growth, development, or fertility of mice. Routine genotyping for detection of *Emx1^IREScre^* and *EPO^flox^* alleles was performed by extraction of genomic DNA collected from ear biopsies upon weaning. Detailed protocol for PCR genotyping is available on request. RNAscope in-situ hybridization was performed on the tissues collected from *NexCre^+/+^::EPOR^fl/fl^* (EPOR WT) [16] *and NexCre^+/-^::EPOR^fl/fl^* (EPOR KO) mouse lines. Immunohistological characterization of EphB4 was performed on NexCre^+/-^::EPOR^+/+^::EphB4^+/+^ (EPOR WT), NexCre^+/-^::EPOR^fl/fl^ (EPOR KO), NexCre^+/-^::EPOR^+/+^::EphB4^+/+^ (EPHB4 WT) and NexCre^+/-^::EPHB4^fl/fl^ (EPHB4 KO) mouse lines.

### mRNA extraction and real-time quantitative reverse transcription polymerase chain reaction (qPCR)

For the characterization of gene expression, KO and WT mice (8-10 weeks old, males and females) were sacrificed by cervical dislocation, followed by isolation of brain (cortex, hippocampus, and cerebellum) and kidneys. Tissue was directly frozen in liquid nitrogen. Total RNA was extracted from tissues by using QIAzol (Qiagen) and miRNeasy Mini Kit (Qiagen, Hilden, Germany). The cDNA reaction mixture (20µl) comprised SuperScript® III Reverse Transcriptase (Thermo Fisher Scientific Life Technologies GmbH, Darmstadt, Germany), Random Hexamer Primer and 1µg of RNA with oligo (dT). Synthesis was performed according to the manufacturer’s instructions. For qPCR reaction mixture, Power SYBR Green PCR Master Mix (Thermo Fisher Scientific Life Technologies) (5µl), cDNA (4µl) and primers (1pmol) were used. The qPCR was performed as described in detail earlier [7, 17]. For Ephrin type-B receptor 4 (*EphB4*) [18], erythropoietin receptor (*EPOR*) [17], erythropoietin (*EPO*) [19], beta-actin (*Actß*) and hypoxanthine guanine phosphoribosyl transferase (*Hprt1*) the following primers were used:

*Hprt1* forward primer: 5‘-GCTTGCTGGTGAAAAGGACCTCTCGAAG-3’

*Hprt1* reverse primer: 5‘-CCCTGAAGTACTCATTATAGTCAAGGGCAT-3’

*ß-actin* forward primer: 5‘-CTTCCTCCCTGGAGAAGAGC-3’

*ß-actin* reverse primer: 5‘-ATGCCACAGGATTCCATACC-3’

*Epo* forward primer: 5‘-CATCTGCGACAGTCGAGTTCTG-3’

*Epo* reverse primer: 5‘-CACAACCCATCGTGACATTTTC-3’

*EpoR* forward primer: 5 ‘CCTCATCTCGTTGTTGCTGA 3’

*EpoR* reverse primer: 5‘ CAGGCCAGATCTTCTGCTG 3’

*EphB4* forward primer: 5 ‘ AGTGGCTTCGAGCCATCAAGA 3’

*EphB4* reverse primer: 5‘ CTCCTGGCTTAGCTTGGGACTTC 3’

The qPCR reactions in 3 technical replicates were performed on LightCycler® 480 System (Roche, Mannheim, Germany). Relative difference in mRNA expression was analyzed by using ΔΔCt method and expression values were normalized to the mean of housekeeping genes, Hprt1 and ß-actin [7].

### Behavioral phenotyping

The role of EPO in forebrain was assessed by performing a basic behavioral battery including elevated plus maze, open field test, Morris water maze, hurdle test, puzzle box [20–25] and higher order cognition by our IntelliCage paradigm [26]. Female *EmxEPO* KO (N=16) and WT (N=16) were used from the age of 7 weeks and tests were performed during the light phase. To avoid stress and anxiety, all experimental mice were habituated before the start of each behavioral test. Group size was based on prior knowledge and technical limitations of behavioral tests following the RRR principle. IntelliCage and Morris water maze have smaller sample sizes due to technical exclusions during testing or analysis (e.g. floating behavior, inability to perform properly, not drinking enough in IntelliCages).

### Magnetic resonance imaging (MRI)

Following anesthesia induction with ketamine (60mg/kg body weight) and medetomidine (0.4mg/kg body weight), the mice were intubated and maintained under 1.5% isoflurane using active ventilation (Animal-Respirator-Advanced^TM^, TSE-Systems). During MRI, each mouse was positioned in a prone orientation with its head fixed using a teeth and palate holder [27]. All MR measurements were performed at a magnetic field strength of 9.4 T (Biospec®, Bruker BioSpin MRI, Ettlingen, Germany) employing the following imaging methods and acquisition parameters: high-resolution T2-weighted images (2D TURBO RARE, TE/TR=55/6000ms, 8 echoes, spatial resolution 40 x 40 x 300µm^3^), and magnetization-transfer (MT) weighted images for volumetric analyses (3D rf-spoiled fast low angle shot (FLASH), TE/TR = 3.4/15.2ms, flip angel 5°, Gaussian-shaped off resonance pulse (off-resonance frequency 3000Hz, power 6µT), spatial resolution 100µm isotropic).

### MRI Data analyses

*Volumetry:* MT-weighted images were first converted to NIfTI and preprocessed through denoising and bias field correction [28] in order to create an unbiased anatomical population template using the python pipeline *twolevel_ants_dbm* (https://github.com/CoBrALab/). In order to quantify the volume of selected brain regions, regions of interest (ROIs) including the lateral ventricles, cerebrum (without ventricles and olfactory bulb), hippocampus, and cerebellum were determined on the study template by manual segmentation using the free, open-source software ITK-SNAP (version 4.0.2). ROIs were than retransformed into the subject space, individually inspected, and, if required, manually corrected. Finally, the respective volume information was extracted.

### RNAscope *in situ* hybridization (ISH)

RNAscope® 2.5 HD Brown Reagent Kit (Cat No. 322300), Advanced Cell Diagnostics (ACD), Hayward, CA, USA was used for the detection of *EphB4* mRNA. ISH was performed as described previously [5] with minor modifications. Briefly, coronal cryosections of 15µm thickness were mounted on SuperFrost Plus Slides, dried and stored at −80°C. Sections were then pretreated by dropwise addition of hydrogen peroxide and incubated for 10min at room temperature (RT). Slides were immersed in boiling target retrieval buffer for 15min, followed by incubation with protease plus for 30min at 40°C. Sections were then hybridized with the corresponding target probe Mm-Ephb4-N-XHs (Cat No. 498201) for 2h at 40°C, followed by a series of amplification and washing steps. Chromogenic signal detection was performed with 3,3’-Diaminobenzidine (DAB) incubation for 20min at RT. Sections were counterstained with 50% Mayer’s hemalum (Merck) and mounted with EcoMount (BioCare Medical). Brown punctate dots were counted in the CA1 region of the hippocampus from one section per mouse (n=5) using a light microscope (Olympus BX-50, Tokyo, Japan) equipped with a 100x oil immersion objective (NA=1.35) and normalized to the area of the respective region (mm²). The quantification was normalized and presented as a percentage calculated to the mean dot number of each ISH experiment [% value = (number of dots/mm²)/(mean dot number/mm²) *100].

### Immunohistochemistry (IHC)

Mice were deeply anesthetized by i.p. injection of Avertin and transcardially perfused via left cardiac ventricle with Ringer’s solution followed by 4% paraformaldehyde (PFA) in sodium phosphate-buffered saline (PBS) 0.1 M, pH 7.4. Dissected brains were post-fixed in 4% formaldehyde and equilibrated subsequently in 30% sucrose dissolved in PBS at 4°C overnight. Brains were then embedded in cryoprotectant (O.C.T.^TM^ Tissue-Tek, Sakura) and stored at −80°C. Whole mouse brains were cut into 30μm thick coronal sections with a Leica CM1950 cryostat (Leica Microsystems, Wetzlar, Germany) and stored at −20°C in 25% ethylene glycol and 25% glycerol in PBS until use. Following blocking and permeabilization with 5% normal horse serum (NHS) in 0.3% Triton X-100 in PBS (PBST) for 1h at RT, primary antibodies were incubated in 5% NHS with 0.3% PBST over 2-3 nights at 4°C. The following primary antibodies were used in this study: anti-EphB4 (Goat, 1:50, AF446; R&D) and anti-CTIP2 (Guinea Pig, 1:500, 325005; Synaptic Systems). After washing, sections were incubated for 2h at room temperature with different secondary antibody cocktails diluted in 3% NHS with 0.3% PBST. The following fluorescently conjugated secondary antibodies were used: donkey anti-goat Alexa Fluor-555 (1:500, A21432; Life Technologies) and donkey anti-Guinea Pig Alexa Fluor-488 (1:500, 706-545-148; Jackson Immunoresearch). Nuclei were stained for 10min at RT with 4′,6-diamidino-2-phenylindole (DAPI; 1:5000; Millipore-Sigma) in PBS. Finally, sections were washed in PBS 0.1 M and mounted on SuperFrostPlus Slides (ThermoFisher) with Aqua-Poly/Mount (Polysciences, Inc).

### Microscope imaging and analysis

For EphB4 and CTIP2 quantification, sections of hippocampus were acquired as tile scans on a confocal laser scanning microscope (LSM 880, Zeiss), furnished with a 40x oil objective (40x/1.4 Oil DIC M27). Quantifications and image processing were performed with FIJI-ImageJ software [29]. EphB4+ CTIP2+ cells were manually counted. CA1 region was determined through manual segmentation. Cell counts were normalized to CA1. Data obtained from 2 to 3 hippocampi/mouse was averaged.

### Single-nuclei RNA sequencing and data processing

Single-cell FASTQ files were aligned to the mm10 mouse genome using 10x Genomics CellRanger count (v6.1.1) to generate gene/cell count matrices. The genome references and gene transfer format (GTF) files were sourced from Ensembl [30] and prepared with “mkref” function, which is available through CellRanger. Alignment was conducted with standard parameters as described in the developer’s manual. To mitigate potential batch effects and address differential gene expression (DGE), background RNA was removed using CellBender (0.2.1106) [31]. Quality control of the alignment and data matrices was performed using downstream processing tools from CellRanger. Filtering, normalization, and cell-types clustering were carried out using Seurat (v4.1.1) [32], which is implemented in R (v4.1.0) [33]. Plausible doublets were removed using the “DoubletFinder” [34]. Cells were filtered based on specific criteria, and normalization was conducted, followed by regression to eliminate the impact of counts mapping to mitochondrial genes. Cells exhibiting high mitochondrial counts (>0.5%) were excluded from subsequent analysis. Clustering of cell types was performed using the top 2000 variable genes expressed across all samples, with a resolution set to 0.6. The construction of the shared-nearest neighbor (SNN) graph and the generation of 2-dimensional embedding for data visualization was carried out using the first 30 PCA dimensions. Cell clusters were assigned into 11 major cell types based on the known transcriptional markers from a literature survey.

### Analyses of EPO/placebo and hypoxia datasets

We leveraged, processed and analyzed single-nuclei RNA sequencing (snRNA-seq) data from our previous EPO study (EPO dataset; GSE220522 [8]) and single-cell RNA sequencing (scRNA-seq) data from our hypoxia dataset (GSE162079). The EPO dataset comprises a cellular map of the mouse hippocampus, encompassing ∼200,000 nuclei from 23 individuals (N = 11 EPO, N = 12 placebo), grouped into 11 major cellular lineages [8]. Feature plots illustrating the expression density of *Emx1*, *EPO*, and *EPOR* were generated on this data using the Nebulosa package in R [35]. To focus on pyramidal neurons, we specifically selected newly formed and mature CA1 neuronal subsets as annotated in the original dataset. For *EPOR* enrichment analysis, further categorized the subsetted data into *EPO*-positive (≥1 transcript count) and *EPO*-negative (0 transcript count) groups. This classification yielded 467 *EPO*-positive nuclei and 19,111 *EPO*-negative nuclei. Genome-wide differential expression analysis was performed using the bimodal test, with false discovery rate (FDR) correction applied via the Benjamini-Hochberg method. We used WebGestalt v. 2019 [36], to identify enriched ontology terms using over-representation analysis (ORA) for phenotypes and transcription factor binding enrichment that were assessed and shown. To investigate *EPOR* expression, we calculated the fraction of cells expressing or not expressing the *EPOR* gene within the defined subsets. The enrichment ratio for each cell type was computed as the ratio of observed to expected values and tested for differential proportions. Statistical significance was determined using a two-way Fisher’s exact test, followed by Bonferroni correction for multiple comparisons. Custom scripts and code used for data processing and visualization are available upon request and will be made publicly accessible on GitHub in a forthcoming update.

### Statistical Analysis

Statistical analyses and bar graphs for behavioral, qPCR and immunohistochemical data was generated using GraphPad Prism-10, R [37] for data analysis, and Grubb’s test to calculate statistical outliers. Results are shown as mean ± standard error of mean (SEM), unless otherwise stated. N number represent number of mice per group. Normal distribution of the data was assessed and statistical tests including unpaired student *t*-tests or repeated measures 2-way ANOVA with *post-hoc* Bonferroni-corrected multiple testing were applied accordingly. P values < 0.05 were considered statistically significant.

## RESULTS

### Consequences of forebrain-specific *EPO* KO on EPO expression

Conditional knockout of *EPO* in forebrain using the *Emx1* promotor leads to a dramatic reduction of *EPO* mRNA expression in hippocampus and cortex, but does – expectedly-not affect the cerebellum which lacks *Emx1* expression [15, 38, 39]. A borderline significant reduction of *EPO* mRNA is also seen in the kidney where *Emx1* is known to be expressed [40] (**Figure 1B**).

### Upregulation of *EPOR and EphB4* upon forebrain-specific *EPO* KO

In tissues that show a remarkable reduction of *EPO* expression, namely hippocampus, cortex and kidney, *EPOR* mRNA displays an upregulation. In contrast, the alternative EPO receptor, EphB4, is increased only in hippocampus, not in cortex and kidney, but additionally in the cerebellum of *EPO* KO mice (**Figure 1 C-D**). These data suggest an apparently compensatory upregulation of specific EPO binding sites in situations where the local ligand production is reduced.

### *EphB4* in brain and its interrelation with *EPOR* expression

Next, *in situ* hybridization and immunohistochemistry were performed in order to elucidate the location of EphB4, the novel EPO receptor in brain. Using NexCre^+/+^::EPOR^fl/fl^ (*EPOR* WT) and NexCre^+/-^::EPOR^fl/fl^ (*EPOR* KO) mice, we could document that *EphB4* mRNA is expressed in pyramidal neurons and enhanced upon *EPOR* deletion in these cells (**Figure 2A-C**). The same holds true for EphB4 protein (**Figure 2 D-F**).

**Figure 2:**
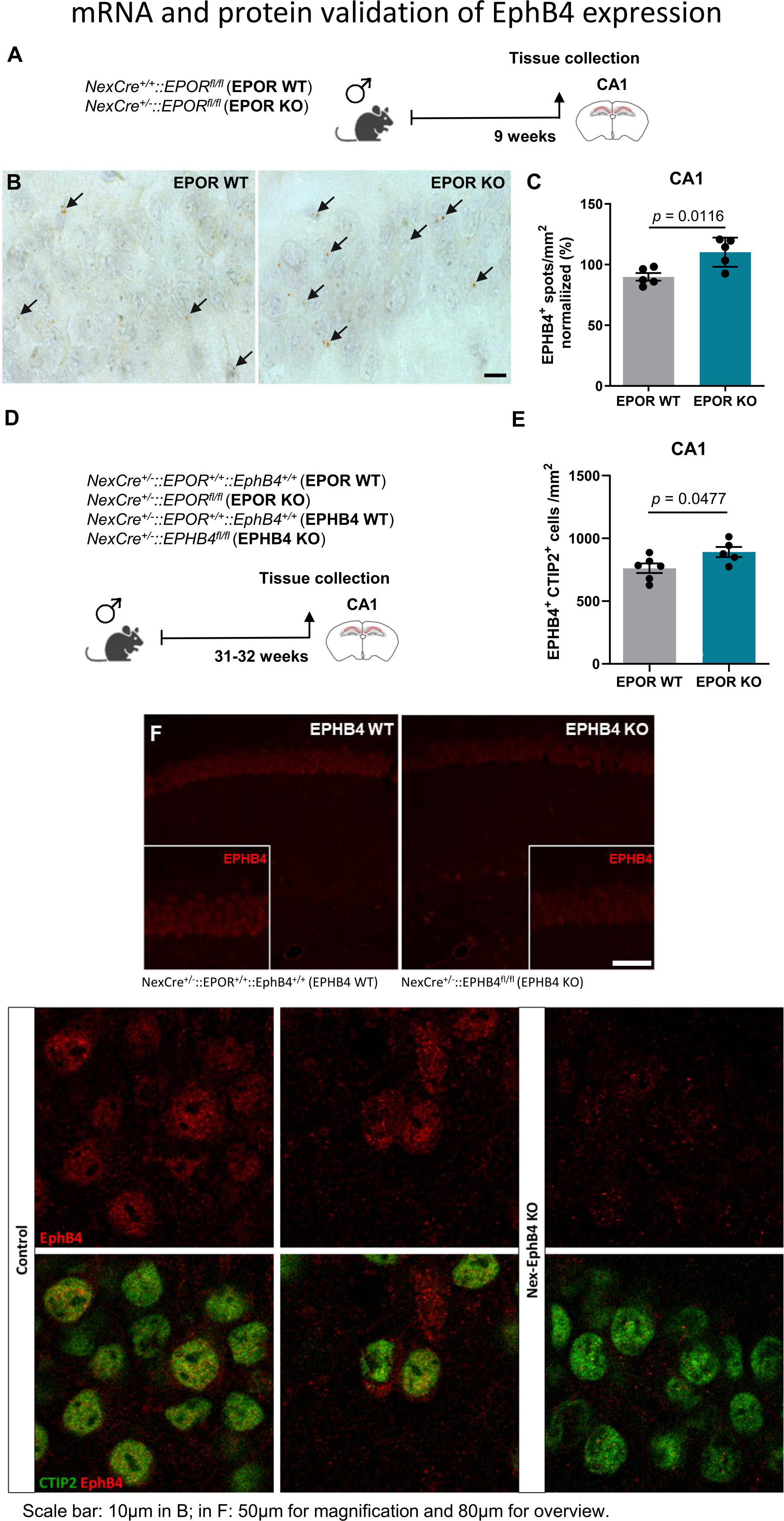
EphB4 expression in the hippocampus and first hints of a regulatory interrelation with EPOR. **(A)** Experimental scheme used for *NexCre^+/+^::EPOR^fl/fl^* (*EPOR* WT) and *NexCre^+/-^::EPOR^fl/fl^* (*EPOR* KO) mice. **(B)** Representative *in situ* hybridization images of the CA1 region from *EPOR* WT and *EPOR* KO mice, demonstrating *EphB4* mRNA expression (brown spots, denoted with arrows) in neurons of the pyramidal layer. **(C)** Quantification of *EphB4* mRNA spots in the CA1 region of *EPOR* WT and *EPOR* KO mice. **(D)** Scheme of experimental timeline and area of analysis (CA1 region) used in *NexCre^+/-^::EPOR^+/+^::EphB4^+/+^* (*EPOR* WT), *NexCre^+/-^::EPOR^fl/fl^* (*EPOR* KO), *NexCre+/-::EPOR^+/+^::EphB4^+/+^* (*EPHB4* WT) and *NexCre^+/-^::EPHB4^fl/fl^* (*EPHB4* KO) mice for immunohistochemical evaluation. **(E)** Quantification of EPHB4+CTIP2+ cells in the pyramidal layer of *EPOR* WT and *EPOR* KO mice. **(F)** Representative images of EPHB4 receptor staining (red) in *EPHB4* WT and *EPHB4* KO mice showing lack of specific immunoreactivity in KO. Unpaired two-tailed Student’s t-test in C & E; scale bar in B: 10µm; in F: 50µm for magnification and 80µm for overview.

### Comprehensive behavioral analysis on forebrain-specific *EPO* KO

Health status and body weight of KO versus WT mice were comparable. In order to see whether reduction of forebrain *EPO* expression would impact any behavioral readouts, we conducted a wide-ranging behavioral battery with female mice. This included anxiety-related tests, evaluation of locomotor and exploratory activity, open field and hurdle test, puzzle box and Morris water maze. As presented in Table 1, there were no appreciable differences between genotypes. It can therefore be concluded that KO mice display an overall normal behavioral phenotype.

### IntelliCage testing reveals superiority of forebrain-specific *EPO* KO

Finally, mice underwent our extensive IntelliCage paradigm (**Figure 3A-B**; **Table 1**), described in great detail elsewhere [26]. Neither the spatial learning and memory nor the episodic-like memory items of the task revealed any appreciable differences in performance between mutants and WT. However, in the most difficult IntelliCage task, namely working memory with its clockwise and counterclockwise challenges, mice with forebrain-specific *EPO* KO consistently demonstrated superior performance. **Magnetic resonance imaging (MRI)** delivered no evidence of altered brain dimensions in forebrain-specific *EPO* KO mice (**Figure 3 C**). Qualitative assessment of the T2-weighted images revealed no observable differences between *EmxEPO* (n=8) and WT (n=8) mice. Likewise, quantitative analysis of brain volumes showed no statistically significant differences between these groups (all p values >0,1).

**Figure 3:**
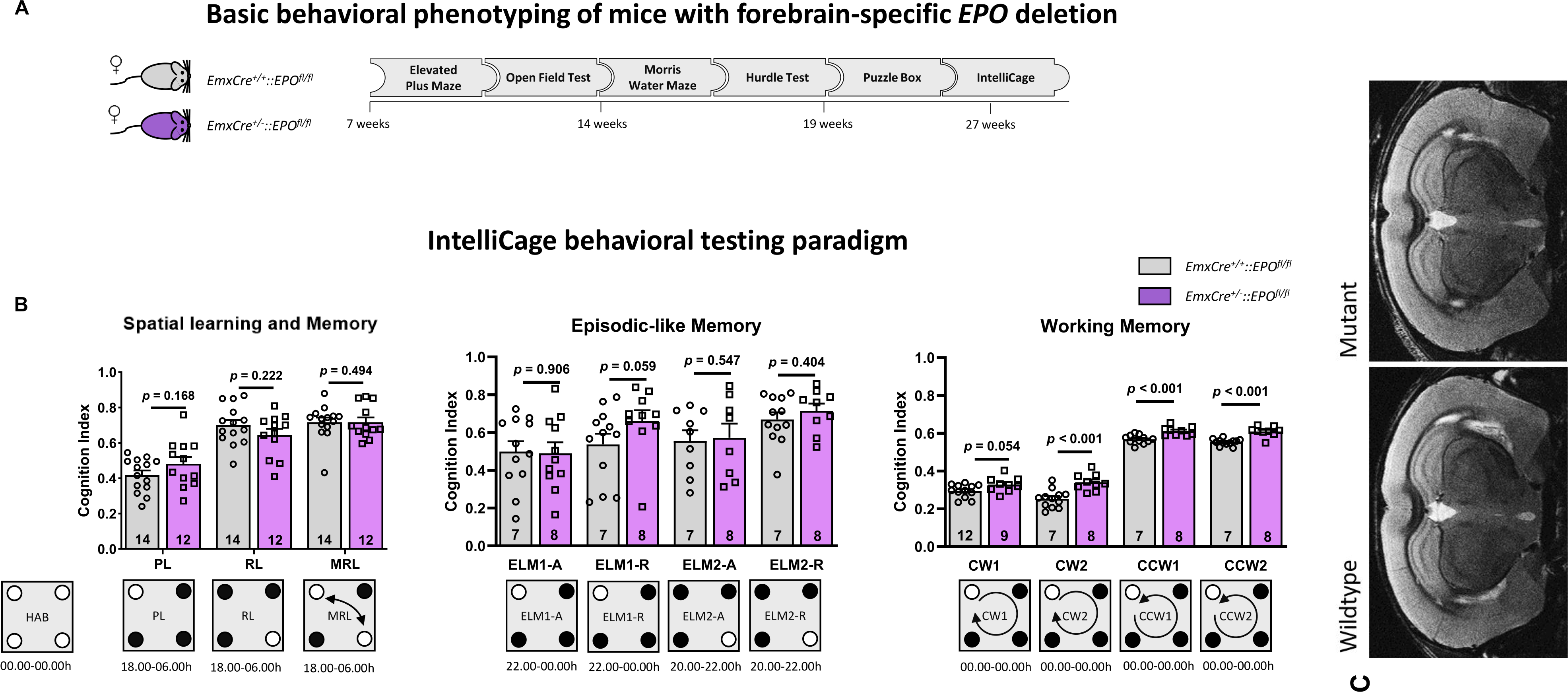
Behavioral phenotyping of mice with forebrain-specific *EPO* deletion. **(A)** Basic behavioral testing. Schematic displays the outline of behavioral experiments performed on *EmxEPO* mice, starting at the age of 7 weeks and concluded at the age of 27 weeks. Detailed behavioral results are presented in **Table1**. **(B)** Higher order cognition was evaluated in our IntelliCage behavioral testing paradigm. Cognition index results for spatial learning and memory, episodic-like memory and working memory are given in scatter bar graphs. A graphical plan of the respective IntelliCage paradigm is presented underneath the bar graphs; HAB: Habituation; PL: Place Learning; RL: Reversal Learning; MRL: Multiple Reversal Learning; ELM: Episodic-like memory; CW: Clockwise; CCW: Counterclockwise; N numbers given in bars; mean ± SEM presented; t-test with Welch’s correction and Mann-Whitney U test were used for statistical analyses. **(C)** MRI: Matched sample sections demonstrate in an exemplary fashion that there was no difference between WT and *EPO* KO mutants.

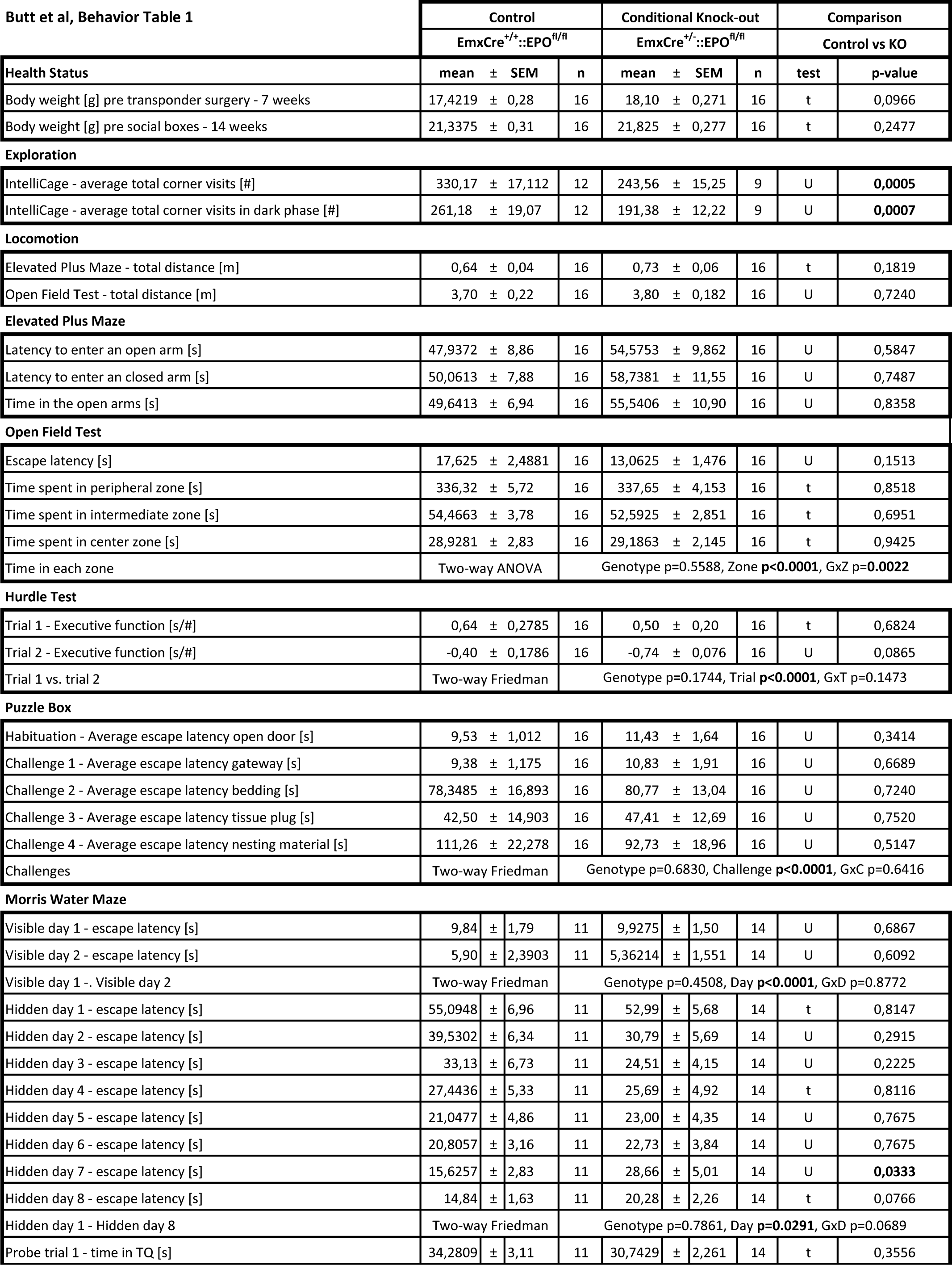
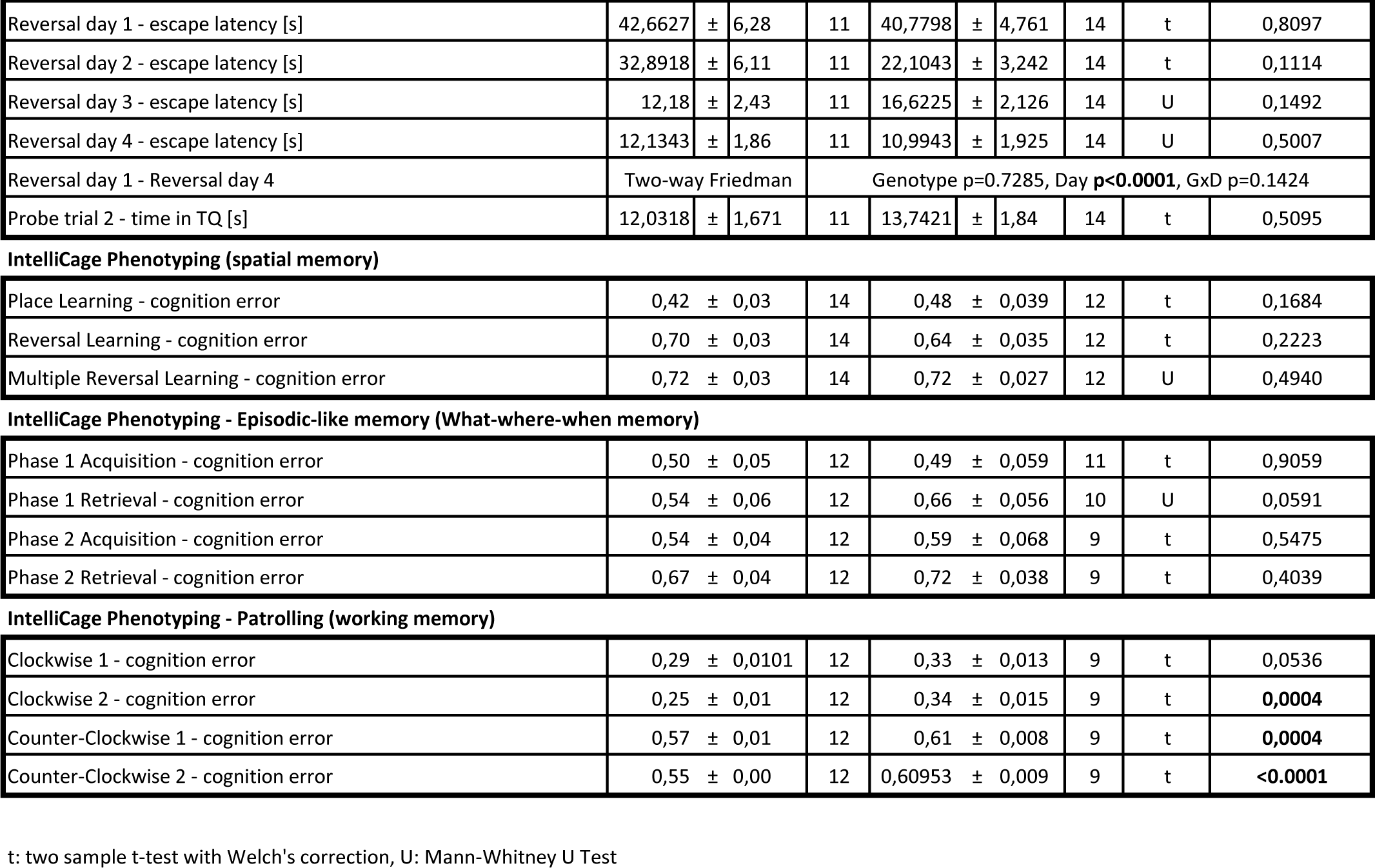

| Butt et al, Behavior Table 1 | Control |  | Conditional Knock-out |  | Comparison |  |
| --- | --- | --- | --- | --- | --- | --- |
|  | EmxCre<sup>+/+</sup>::EPO<sup>fl/fl</sup> |  | EmxCre<sup>+/-</sup>::EPO<sup>fl/fl</sup> |  | Control vs KO |  |
| Health Status | mean ± SEM | n | mean ± SEM | n | test | p-value |
| Body weight [g] pre transponder surgery - 7 weeks | 17,4219 ± 0,28 | 16 | 18,10 ± 0,271 | 16 | t | 0,0966 |
| Body weight [g] pre social boxes - 14 weeks | 21,3375 ± 0,31 | 16 | 21,825 ± 0,277 | 16 | t | 0,2477 |
| Exploration |  |  |  |  |  |  |
| IntelliCage - average total corner visits [#] | 330,17 ± 17,112 | 12 | 243,56 ± 15,25 | 9 | U | 0,0005 |
| IntelliCage - average total corner visits in dark phase [#] | 261,18 ± 19,07 | 12 | 191,38 ± 12,22 | 9 | U | 0,0007 |
| Locomotion |  |  |  |  |  |  |
| Elevated Plus Maze - total distance [m] | 0,64 ± 0,04 | 16 | 0,73 ± 0,06 | 16 | t | 0,1819 |
| Open Field Test - total distance [m] | 3,70 ± 0,22 | 16 | 3,80 ± 0,182 | 16 | U | 0,7240 |
| Elevated Plus Maze |  |  |  |  |  |  |
| Latency to enter an open arm [s] | 47,9372 ± 8,86 | 16 | 54,5753 ± 9,862 | 16 | U | 0,5847 |
| Latency to enter an closed arm [s] | 50,0613 ± 7,88 | 16 | 58,7381 ± 11,55 | 16 | U | 0,7487 |
| Time in the open arms [s] | 49,6413 ± 6,94 | 16 | 55,5406 ± 10,90 | 16 | U | 0,8358 |
| Open Field Test |  |  |  |  |  |  |
| Escape latency [s] | 17,625 ± 2,4881 | 16 | 13,0625 ± 1,476 | 16 | U | 0,1513 |
| Time spent in peripheral zone [s] | 336,32 ± 5,72 | 16 | 337,65 ± 4,153 | 16 | t | 0,8518 |
| Time spent in intermediate zone [s] | 54,4663 ± 3,78 | 16 | 52,5925 ± 2,851 | 16 | t | 0,6951 |
| Time spent in center zone [s] | 28,9281 ± 2,83 | 16 | 29,1863 ± 2,145 | 16 | t | 0,9425 |
| Time in each zone | Two-way ANOVA |  | Genotype p=0.5588, Zone p<0.0001, GxZ p=0.0022 |  |  |  |
| Hurdle Test |  |  |  |  |  |  |
| Trial 1 - Executive function [s/#] | 0,64 ± 0,2785 | 16 | 0,50 ± 0,20 | 16 | t | 0,6824 |
| Trial 2 - Executive function [s/#] | -0,40 ± 0,1786 | 16 | -0,74 ± 0,076 | 16 | U | 0,0865 |
| Trial 1 vs. trial 2 | Two-way Friedman |  | Genotype p=0.1744, Trial p<0.0001, GxT p=0.1473 |  |  |  |
| Puzzle Box |  |  |  |  |  |  |
| Habituation - Average escape latency open door [s] | 9,53 ± 1,012 | 16 | 11,43 ± 1,64 | 16 | U | 0,3414 |
| Challenge 1 - Average escape latency gateway [s] | 9,38 ± 1,175 | 16 | 10,83 ± 1,91 | 16 | U | 0,6689 |
| Challenge 2 - Average escape latency bedding [s] | 78,3485 ± 16,893 | 16 | 80,77 ± 13,04 | 16 | U | 0,7240 |
| Challenge 3 - Average escape latency tissue plug [s] | 42,50 ± 14,903 | 16 | 47,41 ± 12,69 | 16 | U | 0,7520 |
| Challenge 4 - Average escape latency nesting material [s] | 111,26 ± 22,278 | 16 | 92,73 ± 18,96 | 16 | U | 0,5147 |
| Challenges | Two-way Friedman |  | Genotype p=0.6830, Challenge p<0.0001, GxC p=0.6416 |  |  |  |
| Morris Water Maze |  |  |  |  |  |  |
| Visible day 1 - escape latency [s] | 9,84 ± 1,79 | 11 | 9,9275 ± 1,50 | 14 | U | 0,6867 |
| Visible day 2 - escape latency [s] | 5,90 ± 2,3903 | 11 | 5,36214 ± 1,551 | 14 | U | 0,6092 |
| Visible day 1 -. Visible day 2 | Two-way Friedman |  | Genotype p=0.4508, Day p<0.0001, GxD p=0.8772 |  |  |  |
| Hidden day 1 - escape latency [s] | 55,0948 ± 6,96 | 11 | 52,99 ± 5,68 | 14 | t | 0,8147 |
| Hidden day 2 - escape latency [s] | 39,5302 ± 6,34 | 11 | 30,79 ± 5,69 | 14 | U | 0,2915 |
| Hidden day 3 - escape latency [s] | 33,13 ± 6,73 | 11 | 24,51 ± 4,15 | 14 | U | 0,2225 |
| Hidden day 4 - escape latency [s] | 27,4436 ± 5,33 | 11 | 25,69 ± 4,92 | 14 | t | 0,8116 |
| Hidden day 5 - escape latency [s] | 21,0477 ± 4,86 | 11 | 23,00 ± 4,35 | 14 | U | 0,7675 |
| Hidden day 6 - escape latency [s] | 20,8057 ± 3,16 | 11 | 22,73 ± 3,84 | 14 | U | 0,7675 |
| Hidden day 7 - escape latency [s] | 15,6257 ± 2,83 | 11 | 28,66 ± 5,01 | 14 | U | 0,0333 |
| Hidden day 8 - escape latency [s] | 14,84 ± 1,63 | 11 | 20,28 ± 2,26 | 14 | t | 0,0766 |
| Hidden day 1 - Hidden day 8 | Two-way Friedman |  | Genotype p=0.7861, Day p=0.0291, GxD p=0.0689 |  |  |  |
| Probe trial 1 - time in TQ [s] | 34,2809 ± 3,11 | 11 | 30,7429 ± 2,261 | 14 | t | 0,3556 |

|  |  |  |  |  |  |  |  |  |  |  |
| --- | --- | --- | --- | --- | --- | --- | --- | --- | --- | --- |
| Reversal day 1 - escape latency [s] | 42,6627 | ± | 6,28 | 11 | 40,7798 | ± | 4,761 | 14 | t | 0,8097 |
| Reversal day 2 - escape latency [s] | 32,8918 | ± | 6,11 | 11 | 22,1043 | ± | 3,242 | 14 | t | 0,1114 |
| Reversal day 3 - escape latency [s] | 12,18 | ± | 2,43 | 11 | 16,6225 | ± | 2,126 | 14 | U | 0,1492 |
| Reversal day 4 - escape latency [s] | 12,1343 | ± | 1,86 | 11 | 10,9943 | ± | 1,925 | 14 | U | 0,5007 |
| Reversal day 1 - Reversal day 4 | Two-way Friedman |  |  | Genotype p=0.7285, Day p<0.0001, GxD p=0.1424 |  |  |  |  |  |  |
| Probe trial 2 - time in TQ [s] | 12,0318 | ± | 1,671 | 11 | 13,7421 | ± | 1,84 | 14 | t | 0,5095 |

IntelliCage Phenotyping (spatial memory)
|  |  |  |  |  |  |  |  |  |  |  |
| --- | --- | --- | --- | --- | --- | --- | --- | --- | --- | --- |
| Place Learning - cognition error | 0,42 | ± | 0,03 | 14 | 0,48 | ± | 0,039 | 12 | t | 0,1684 |
| Reversal Learning - cognition error | 0,70 | ± | 0,03 | 14 | 0,64 | ± | 0,035 | 12 | t | 0,2223 |
| Multiple Reversal Learning - cognition error | 0,72 | ± | 0,03 | 14 | 0,72 | ± | 0,027 | 12 | U | 0,4940 |

IntelliCage Phenotyping - Episodic-like memory (What-where-when memory)
|  |  |  |  |  |  |  |  |  |  |  |
| --- | --- | --- | --- | --- | --- | --- | --- | --- | --- | --- |
| Phase 1 Acquisition - cognition error | 0,50 | ± | 0,05 | 12 | 0,49 | ± | 0,059 | 11 | t | 0,9059 |
| Phase 1 Retrieval - cognition error | 0,54 | ± | 0,06 | 12 | 0,66 | ± | 0,056 | 10 | U | 0,0591 |
| Phase 2 Acquisition - cognition error | 0,54 | ± | 0,04 | 12 | 0,59 | ± | 0,068 | 9 | t | 0,5475 |
| Phase 2 Retrieval - cognition error | 0,67 | ± | 0,04 | 12 | 0,72 | ± | 0,038 | 9 | t | 0,4039 |

IntelliCage Phenotyping - Patrolling (working memory)
|  |  |  |  |  |  |  |  |  |  |  |
| --- | --- | --- | --- | --- | --- | --- | --- | --- | --- | --- |
| Clockwise 1 - cognition error | 0,29 | ± | 0,0101 | 12 | 0,33 | ± | 0,013 | 9 | t | 0,0536 |
| Clockwise 2 - cognition error | 0,25 | ± | 0,01 | 12 | 0,34 | ± | 0,015 | 9 | t | 0,0004 |
| Counter-Clockwise 1 - cognition error | 0,57 | ± | 0,01 | 12 | 0,61 | ± | 0,008 | 9 | t | 0,0004 |
| Counter-Clockwise 2 - cognition error | 0,55 | ± | 0,00 | 12 | 0,60953 | ± | 0,009 | 9 | t | <0.0001 |

t: two sample t-test with Welch's correction, U: Mann-Whitney U Test

### snRNA-seq reveals specific Emx1 expression patterns

In the present work, we employed mice with forebrain-specific deletion of *EPO* under control of the *Emx1* promoter [15]. In this context, we wondered whether *Emx1* expression itself in adult mice would be influenced by EPO. We thus investigated hippocampal transcriptional patterns of 23 mice, 11 treated with rhEPO and 12 with placebo, at the level of individual nuclei to check *Emx1* gene expression. Analyzing approximately 200,000 single nuclei enabled us to uncover an amazing heterogeneity within the hippocampi. Through clustering based on similar transcriptional profiles, we identified 36 distinct clusters, which were further classified into 10 primary lineages and 1 neuroimmune cluster, each exhibiting unique gene expression patterns (**Figure 4A**).

**Figure 4:**
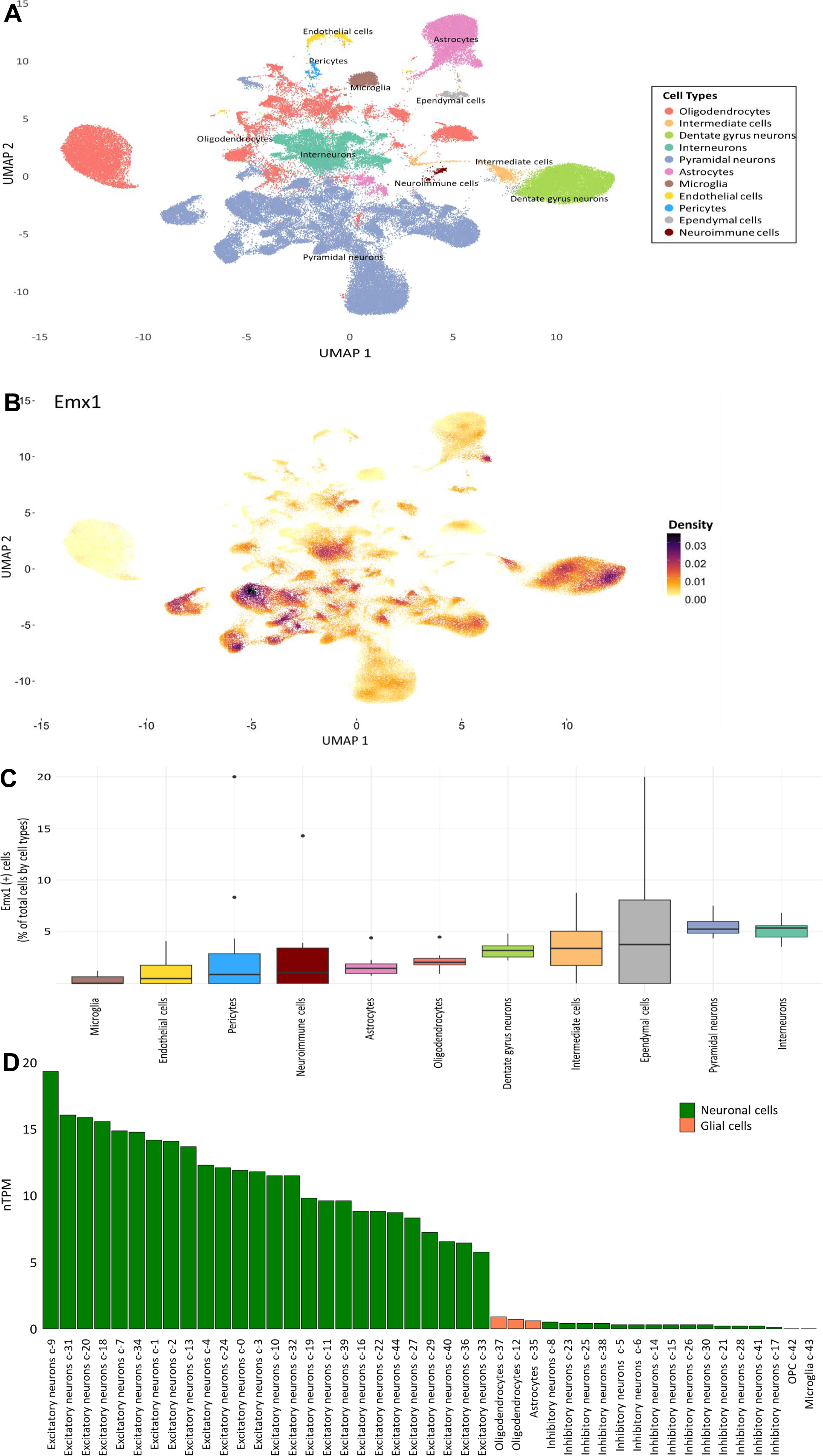
Single-nuclei RNA sequencing (snRNA-seq) allows to determine *Emx1* gene expression in adult hippocampus. **(A)** Two-dimensional Uniform Manifold Approximation Plot (UMAP) from snRNA-seq of merged 12 adult hippocampal samples of mice treated with either rhEPO (N = 6) or placebo (N = 6). **(B)** Density plot illustrating *Emx1* gene expression within snRNA-seq data from the 12 adult hippocampal samples. **(C)** Percentage (%) of *Emx1* positive cells per cell type in the hippocampal snRNA-seq dataset. **(D)** Expression profile of *Emx1* gene within the human brain, sourced from the Human Protein Atlas database [51].

Examining the density of *Emx1* gene expression revealed significant variation among cell clusters (**Figure 4B**). The proportion of *Emx1*-expressing cells (*Emx1+*) was determined using the Percent_Expressing function from scCustomize [41], and the distribution of percentages of *Emx1+* cells across clusters is depicted in **Figure 4C**. Notably, interneurons (with a median of 5.35% *Emx1+* cells and a mean of 5.16%) and pyramidal neurons (with a median of 5.25% *Emx1+* cells and a mean of 5.54%) exhibited the highest percentages of *Emx1+* cells. In contrast, microglia had the lowest percentage of *Emx1+* cells (with a median of 0.00% *Emx1+* cells and a mean of 0.30%). The ependymal cell cluster exhibited the most notable variability in *Emx1+* cell percentages, with a median of 3.76% and a mean of 5.96%, indicating a wide range of expression levels within this specific cell population. Moreover, our analysis did not reveal any significant differential expression of *Emx1* gene between rhEPO and placebo treated groups (log2FC < 0.1).

Furthermore, *Emx1* expression was cross-validated using data from the Human Protein Atlas (**Figure 4D**). In glial cells (including oligodendrocytes, astrocytes, OPCs, and microglia), *Emx1* was found to be expressed at low levels (normalized transcripts per million, nTPM < 1). Conversely, in neuronal cells, it exhibited higher expression levels in excitatory neurons (with an nTPM range of 20 to 6 and mean and median values of approximately 12) while being expressed at lower levels in inhibitory neurons (nTPM < 1).

### Compensatory upregulation validated by snRNA-seq

To delve deeper into *EPOR* expression, we re-analyzed our publicly available snRNA-seq datasets (GSE220522). These datasets encompass the transcriptomes of ∼200,000 nuclei isolated from the hippocampi of 23 mice, treated with rhEPO or placebo[8]. Focusing on *Emx1* expression, we analyzed 11 hippocampal lineages and found its expression essentially restricted to the dentate gyrus (DG) and newly formed pyramidal neurons (**Figure 4**). In line with this result, we narrowed our analysis to pyramidal neuronal lineages from the DG and CA1 regions.

Using this refined dataset, we examined the distribution of *EPOR* expression across 3 newly formed and 2 mature neuronal types as defined in our previous study [8]. Consistent with earlier findings (**Figure 4B**), *Emx1* expression was predominantly enriched in newly formed rather than mature pyramidal clusters (**Figure 5A**). However, *EPO* and *EPOR* exhibited a mosaic expression pattern across both newly formed and mature CA1 neurons (**Figure 5A**).

**Figure 5:**
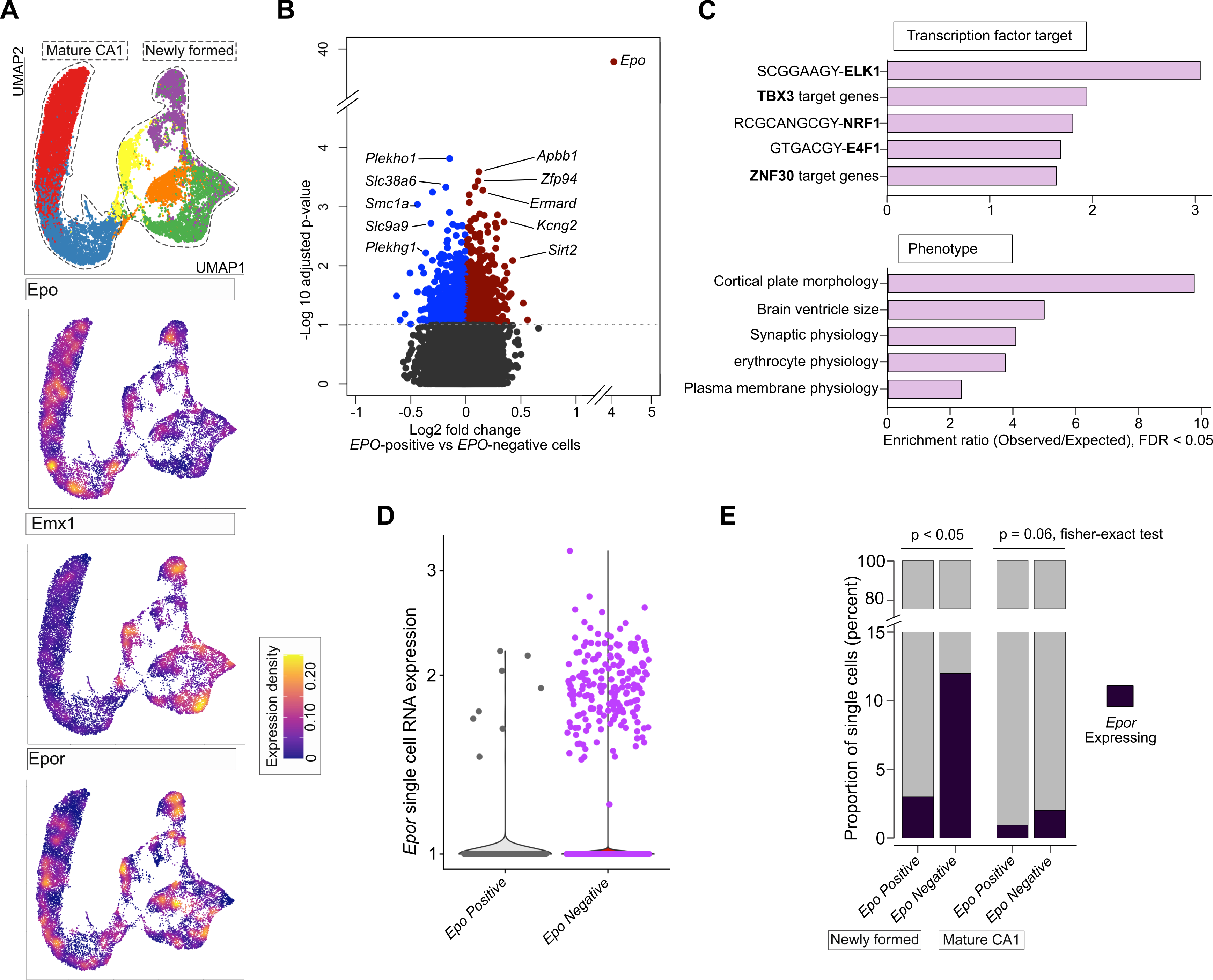
Single-nuclei RNA sequencing (snRNA-seq) for exploring EPO and EPOR expression. **(A)** UMAP clustering of nuclei from the mature CA1 and newly formed clusters are labelled and encircled (top). Feature plot based on the UMAP plot showing the single nuclei expression of *EPO, Emx1 and EPOR* in pyramidal neurons. **(B)** Volcano plot showing genes that are differentially expressed between *EPO*-positive and *EPO*-negative pyramidal neurons. The horizontal dashed line indicates -Log_10_P = 1 (FDR corrected two-way Fisher’s exact test). Differentially expressed genes are colored blue (down-regulated) and red (up-regulated). *The top differentially expressed genes* are shown which contribute to the gene ontology in the next figure subset. **(C)** Enriched pathways, and gene ontologies, correspond to phenotypes and transcription factor target genes among genes that are differentially expressed (colored in the previous plot). Data is shown as an enrichment score (observed/expected), with -log10 hypergeometric P-value < 0.05. **(D)** *EPOR* expression in the analyzed pyramidal cell types is upregulated in EPO-negative (purple) when compared with EPO-positive (grey) nuclei. Every dot shown on the plot represents a nucleus from snRNA-seq data. Transcript abundance in snRNA-Seq data is shown as log-transformed expression of *EPOR* normalized using the transcriptome. **(E)** The stacked bar plot shows the percentage of nuclei expressing EPOR in 2 distinct groups of nuclei from EPO-positive and EPO-negative pyramidal neurons.

To simulate theoretical *EPO* knockout conditions, we categorized pyramidal neurons into *EPO-*positive (≥1 transcript count) and *EPO*-negative (0 transcript count) groups, termed *EPO*-positive and *EPO*-negative cells, respectively. A differential gene expression analysis using a bimodal test revealed 739 significantly differentially expressed genes (adjusted p-value < 0.05) between these groups (**Figure 5B**, Figure 5—Source Data 1). Gene ontology enrichment highlighted phenotypes associated with “cortical plate morphology”, “brain ventricle size”, and “synaptic, plasma membrane, and erythrocyte physiologies” (**Figure 5C**).

Next, we identified transcription factors potentially regulating these genes using WebGestalt analysis [42]. Key candidates included ELK1, NRF1, TBX3, E4F1, and ZNF30 (**Figure 5C**). Notably, these transcription factors are implicated in hippocampal learning, survival and memory: ELK1 [43, 44], NRF1 [45, 46], E4F1 [47, 48], and ZNF30 [49]. These findings, observed in both EPO/placebo and hypoxia datasets, suggest a mechanistic link between *EPO* expression in hippocampal pyramidal neurons and associated phenotypes observed across numerous studies, including the present one.

Pushing further, we assessed *EPOR* expression patterns and the proportion of pyramidal neurons expressing *EPOR* in *EPO*-positive and *EPO*-negative conditions. While *EPOR* RNA levels were elevated in *EPO*-negative cells, the increase was modest (**Figure 5D**). This observation prompted us to test whether the probability of *EPOR* expression differing between these groups could be attributed to the higher number of cells in *EPO*-negative conditions. Thus, we determined the percent *EPOR* expressing cells in both *EPO*-positive and *EPO*-negative cells. Remarkably, *EPO*-negative cells showed a significantly higher number of *EPOR*-expressing cells, particularly among newly formed pyramidal neurons (**Figure 5E**).

Collectively, these findings, based on our snRNA-seq dataset, propose a compensatory mechanism wherein *EPO* loss drives elevated *EPOR* expression, supporting the respective observation in the forebrain of *EmxEPO* mice, and adding another layer to the intricate regulation of EPO signaling in hippocampal pyramidal neurons.

## DISCUSSION

The present work has originally been designed to explore consequences of reduced expression of EPO in the forebrain regarding behavior, cognition and other phenotypical measures. Upon *EmxEPO* KO, the ligand, EPO, is diminished, however, not completely abolished. The ‘leftover expression’ is explained by (1) the presence of cell types expressing EPO but not Emx1, namely oligodendrocyte precursor cells (OPC) and microglia, (2) the availability of peripherally produced EPO, circulating through the blood stream and cerebrospinal fluid. Surprisingly, we find nearly no phenotypical changes under these conditions of blunted EPO expression. This may be explained by the here observed substantial compensatory upregulation of its receptors, EPOR and EphB4, in the forebrain. Exploiting our snRNA-seq dataset, we confirm this novel compensatory regulation within the EPO system and gain further molecular insights into EPO-related phenotypes.

Although 20–30% upregulation of EPOR and EphB4 as observed in heterozygous EmxEPO KO mice of the current study may seem quantitatively modest, such regulatory shifts are likely biologically meaningful in the context of neuroplasticity and receptor sensitivity. In fact, small changes in key signaling pathways can have amplified downstream effects on synaptic remodeling and circuit integration, making these subtle changes biologically/physiologically meaningful. EPOR expression in the brain is naturally extremely low [4]. The increase in classical EPOR - and, in particular, the additional upregulation of the other EPO receptor, EphB4 (having ephrinB2 as its canonical ligand [50]) - can enhance or diversify downstream signaling. Functionally, this molecular compensation, achieved through the upregulation of EPOR and EphB4 in Emx1-Cre EPO knockout mice, aligns with improved performance in the most demanding part of the IntelliCage working memory tasks, indicating that the observed mild receptor upregulation has a boosting effect on cognitive function. A more complete ligand depletion would be expected to be lethal and not to amplify the phenotype. Residual EPO transcripts arise primarily from Emx1 negative cell types. Eliminating these sources would necessitate additional Cre drivers and could provoke secondary adaptations such as enhanced hypoxia-inducible factor (HIF) activity that might again compromise the net signaling. For these reasons, we consider the observed receptor increase and the accompanying behavioral effect a biologically significant demonstration of homeostatic compensation without triggering severe structural or viability deficits.

In fact, exploring our snRNA-seq dataset, we find upregulation of *EPOR* expression in *EPO*-negative cells. This suggests an adaptive response aimed at maintaining homeostasis following loss of *EPO*. The differential expression of transcription factors ELK1 and NRF1 between *EPO*-positive and *EPO*-negative cells offers a potential molecular basis for phenotypes observed in seminal studies demonstrating EPO effects on learning and memory. ELK1 and NRF1, known to regulate immediate early genes, STAT signaling, and hippocampal memory-associated pathways [44] may mediate EPO-induced enhancements in cognitive function. These factors could represent critical nodes through which EPO exerts its effects, linking our present observations to established mechanisms of learning and memory.

While our snRNA-seq data provide valuable insights, they must be interpreted cautiously. The inherent limitations of this method, including dropout effects and its correlative nature, are particularly relevant to analyses of *EPO* and *EPOR* expression. These constraints call for complementary approaches to expand upon our findings. Such essential complementary approach has been applied here by means of *EmxEPO* KO mice.

We acknowledge the limitation that – due to the present lack of respective animal permits - the current study did not evaluate the effects of exogenous EPO (rhEPO) administration or inspiratory hypoxia treatment in *EmxEPO* KO mice on downstream regulation of EPOR or EphB4 and related molecular and cognitive changes.

Nevertheless, the study of *EmxEPO* KO mice, lacking EPO expression in the forebrain, allowed to identify as yet unknown compensatory mechanisms within the EPO system: Lack of EPO is apparently compensated for by upregulation of EPOR and the novel EPO receptor in brain, EphB4, the regulation of which seems interrelated.

## DATA AVAILABILITY

The snRNA-seq data that support the findings of this study are available from the authors upon request. Hypoxia dataset (GSE162079), EPO dataset (GSE220522).

## AUTHOR CONTRIBUTIONS

Supervision: HE

Funding acquisition: HE, KAN

Concept and design: HE, UJB

Data acquisition/generation: UJB, UÇ, YC, LY, SB, MS

Data analyses/interpretation: UJB, AFW, VB, KAN, SB, MS, HE

Drafting the manuscript: HE, UJB

Drafting display items: UJB, AFW, UÇ, HE

Continuous critical input, review & editing: KAN, HE

**All authors read and approved the final version of the manuscript.**

## ACKNOWLEDGEMENTS

This work has been funded by the European Research Council (ERC) Advanced Grant to HE under the European Union’s Horizon Europe research and innovation programme (acronym *BREPOCI;* grant agreement No 101054369). Furthermore, the study has been supported by the Max Planck Society and the Max Planck Förderstiftung. Research in the labs of HE and KAN is sponsored by DFG TRR-274/1 2020-408885537. YC has been recipient of a grant from the Peter and Traudl Engelhorn Foundation. VB has received a grant from the DFG (Project number 513977564). UÇ obtained funding from the IMPRS-Genome Science PhD program. KAN has been supported by the Adelson Medical Research Foundation.

## ANIMAL ETHICS APPROVAL

All methods were performed in accordance with the relevant guidelines and regulations. Approval has been obtained from the local authorities (Animal Care and Use Committee: Niedersächsisches Landesamt für Verbraucherschutz und Lebensmittelsicherheit, LAVES) following the German Animal Protection Law (AZ 33.19-42502-04-18/2803 & AZ 33.19-42502-04-17/2393). All experiments were conducted by investigators unaware of genotypes and group assignment (′fully blinded′).

